# CHiCAGO: Robust Detection of DNA Looping Interactions in Capture Hi-C data

**DOI:** 10.1101/028068

**Authors:** Jonathan Cairns, Paula Freire-Pritchett, Steven W. Wingett, Csilla Várnai, Andrew Dimond, Vincent Plagnol, Daniel Zerbino, Stefan Schoenfelder, Biola-Maria Javierre, Cameron Osborne, Peter Fraser, Mikhail Spivakov

## Abstract

Capture Hi-C (CHi-C) is a state-of-the art method for profiling chromosomal interactions involving targeted regions of interest (such as gene promoters) globally and at high resolution. Signal detection in CHi-C data involves a number of statistical challenges that are not observed when using other Hi-C-like techniques. We present a background model, and algorithms for normalisation and multiple testing that are specifically adapted to CHi-C experiments, in which many spatially dispersed regions are captured, such as in Promoter CHi-C. We implement these procedures in CHiCAGO (http://regulatorygenomicsgroup.org/chicago), an open-source package for robust interaction detection in CHi-C. We validate CHiCAGO by showing that promoter-interacting regions detected with this method are enriched for regulatory features and disease-associated SNPs.

## BACKGROUND

Chromosome Conformation Capture (3C) technology has revolutionised the analysis of nuclear organisation, leading to important insights into gene regulation [1]. While the original 3C protocol tested interactions between a single pair of candidate regions (“one vs one”), subsequent efforts focused on increasing the throughput of this technology (4C, “one vs all”; 5C, “many vs many”), culminating in the development of Hi-C, a method that interrogated the whole nuclear interactome (“all vs all”) [1, 2]. The extremely large number of possible pairwise interactions in Hi-C samples, however, imposes limitations on the realistically achievable sequencing depth at individual interactions, leading to reduced sensitivity. The recently-developed Capture Hi-C (CHi-C) technology uses sequence capture to enrich Hi-C material for multiple genomic regions of interest (hereafter referred to as “baits”), making it possible to profile the global interaction profiles of many thousands of regions globally (“many vs all”) and at a high resolution (**Fig. 1**) [3–7].

**Figure 1:**
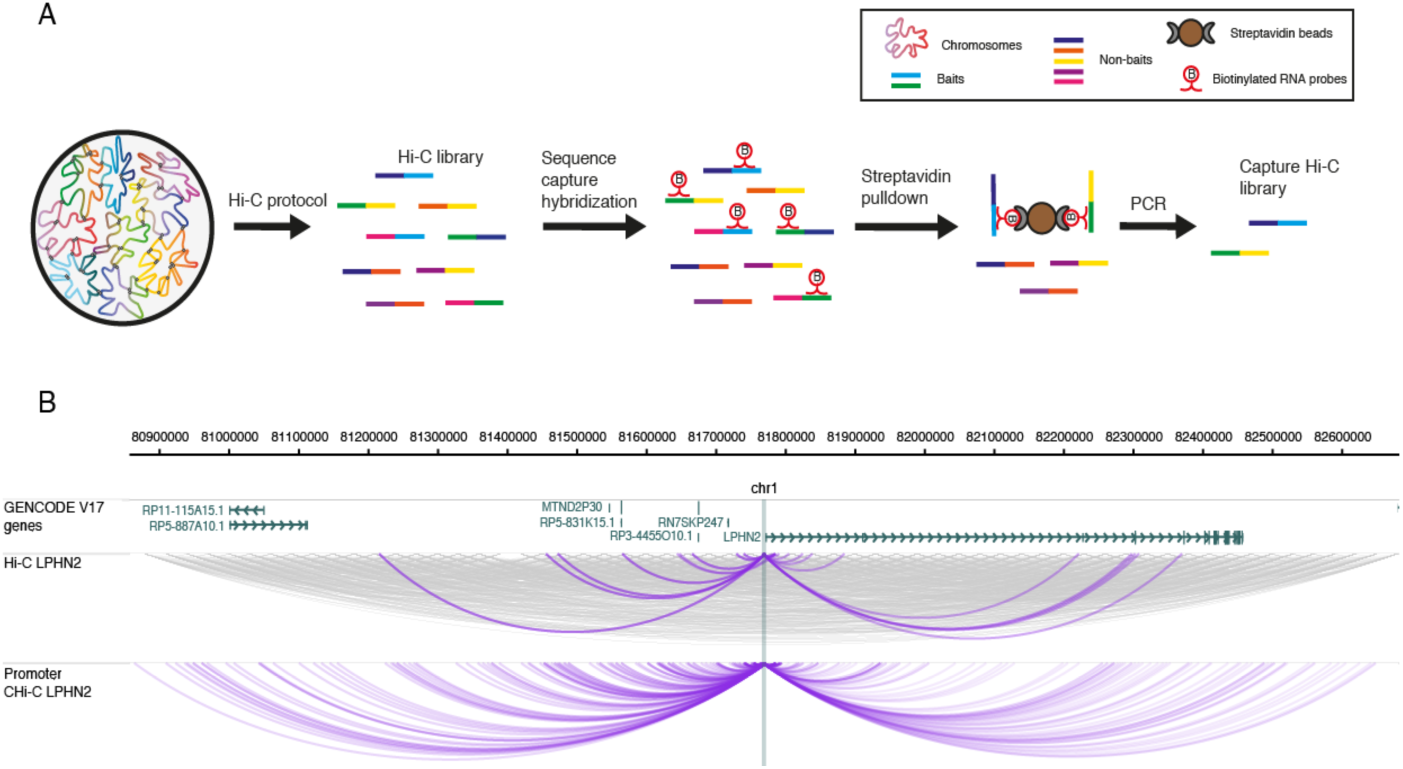
The outline of Capture Hi-C. (A) Outline of the CHi-C protocol. A Hi-C library is hybridized to a capture system that consists of biotinylated RNA probes targeting the ends of DNA restriction fragments. After hybridization, streptavidin pulldown is performed to filter for fragments that have hybridized with the RNA probes, leading to enrichment in baited fragments (“baits”). Following a limited-cycle PCR amplification, the CHi-C library is ready to be analysed by massively parallel paired-end sequencing. (B) The interactome of the *LPHN2* promoter region in GM12878 cells. The top panel shows a 1.8Mb region containing the *LPHN2* gene. The middle panel shows raw read-pairs from the Hi-C library. All read pairs sequenced for these regions are shown in grey. In purple, we show only the read-pairs that contain the *LPHN2* promoter in one of the fragment ends. The bottom panel shows raw read-pairs from the Promoter CHi-C library from [3]. The WashU EpiGenome Browser [16, 17] was used to create this figure.

CHi-C data possess statistical properties that set them apart from other 3C/4C/Hi-C-like methods. First, in contrast to traditional Hi-C or 5C, baits in CHi-C comprise a subset of restriction fragments, while any fragment in the genome can be detected on the “other end” of an interaction. This asymmetry of CHi-C interaction matrices is not accounted for by the normalisation procedures developed for traditional Hi-C and 5C [8–10]. Secondly, CHi-C baits, but not other ends, have an additional source of bias associated with uneven capture efficiency. In addition, the need for detecting interactions globally and at a single-fragment resolution creates specific multiple testing challenges that are less pronounced with binned Hi-C data or the more focused 4C and 5C assays, which involve fewer fragment pairs tested for interaction. Finally, CHi-C designs such as Promoter CHi-C and HiCap [3–5, 11] involve large numbers (many thousands) of spatially dispersed baits. This presents the opportunity to increase the robustness of signal detection by sharing information across baits. Such sharing is impossible in the analysis of 4C data that focuses on only a single bait, and is of limited use in 4C-seq containing a small number of baits [12–14],

These distinct features of CHi-C data have prompted us to develop a bespoke statistical model and a background correction procedure for detecting significant interactions in CHi-C data at a single restriction fragment resolution. The algorithm, termed CHiCAGO (“Capture Hi-C Analysis Of Genomic Organisation”), is presented here and implemented as an open-source R package. CHiCAGO features a novel background correction procedure and a two-component convolution background model accounting for both real, but expected interactions, as well as assay and sequencing artefacts. In addition, CHiCAGO implements a weighted false discovery control procedure that builds on the theoretical foundations of Genovese *et al*. [15]. This procedure specifically accommodates the fact that increasingly larger numbers of tests are performed at regions where progressively smaller numbers of interactions are expected.

We demonstrate the efficacy of CHiCAGO on two datasets: one from the human lymphoblastoid cell line GM12878 [3] (see **Fig. 2** for examples) and another from mouse embryonic stem cells [4]. We further show that CHiCAGO-detected interactions are enriched for regulatory regions and relevant disease-associated SNPs.

**Figure 2:**
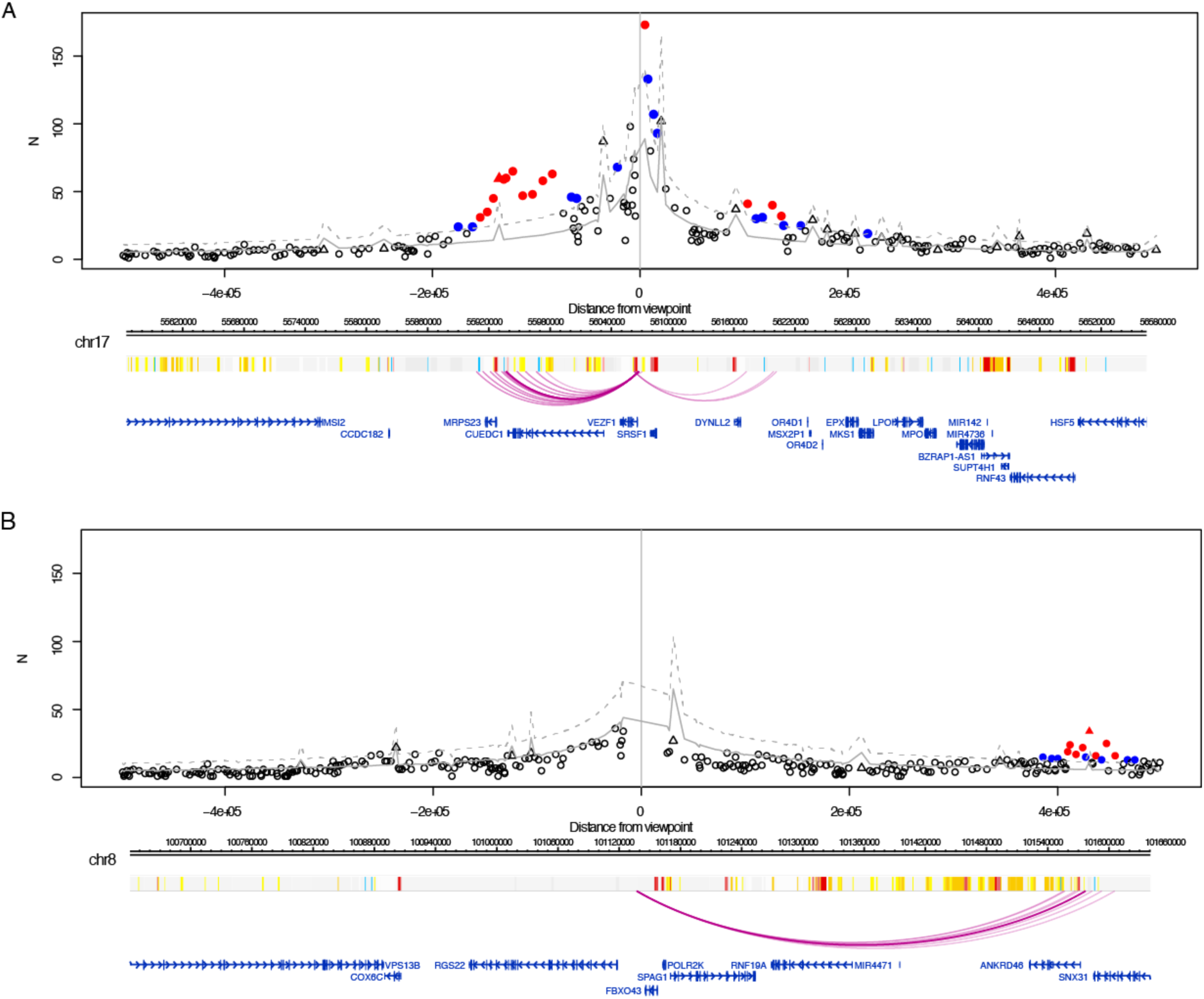
Examples of interactions called by CHiCAGO. Top panels: Plots showing the read counts from bait-other end pairs within 500 kb (upstream and downstream) of two baits, containing the promoters of (A) *VEZF1* and (B) RGS22 in GM12878 cells. Significant interactions detected by CHiCAGO (score >= 5) are shown in red, and sub-threshold interactions (3 <= score <5) are shown in blue. Triangles indicate bait-to-bait interactions. Grey lines show expected counts and dashed lines the upper bound of the 95% confidence intervals. (Note that bait-to-bait interactions have higher expected read counts than bait-to-non-bait interactions spanning the same distance). Bottom panels: the genomic maps of the corresponding regions, with coloured bars showing “chromatin colours” obtained from performing chromatin segmentation with chromHMM [18]: red – active promoter; pink – poised/repressed promoter; orange – strong enhancer; yellow – weak enhancer; blue – insulator.

## RESULTS

### Methodological foundations of CHiCAGO

#### A convolution background model for Hi-C data

The background signals in CHi-C decrease as the genomic distance between the bait and other end increases (**Fig. 3**), as in other 3C/Hi-C-like methods [6–10, 12, 13, 19, 20]. It is generally accepted that this effect reflects the reduction in the frequency of random collisions between genomic fragments owing to constrained Brownian motion of chromatin, in a manner consistent with molecular dynamics simulations [21]. We model the read counts arising from these “Brownian collisions” as a negative binomial random variable whose expected levels are a function of genomic distance with further adjustment for bias resulting from the properties of individual fragments.

**Figure 3:**
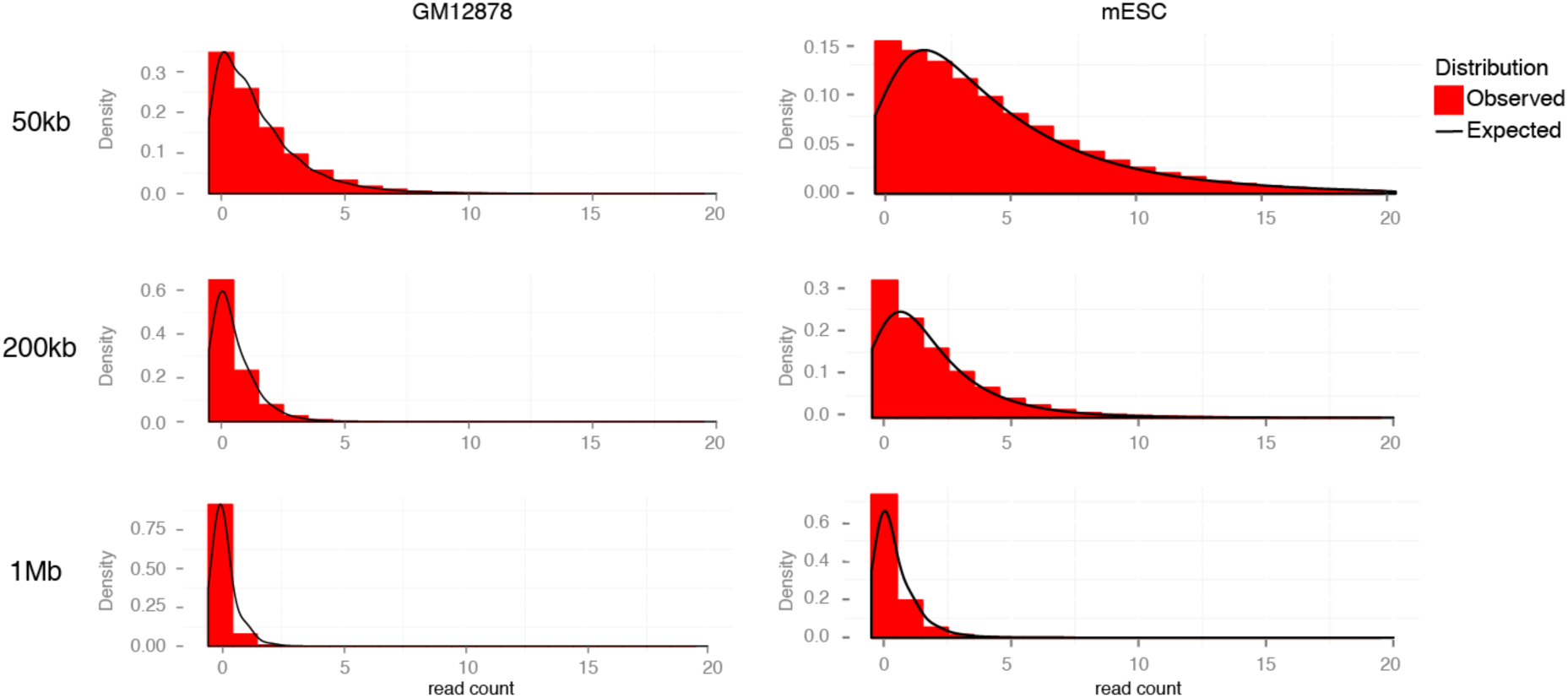
Read count distributions in a typical CHi-C experiment and their fit to the CHiCAGO background model. Histograms showing read count distributions for fragment pairs spanning different distance ranges (*/-20 kb) in a biological replicate of GM12878 (left) and mESC CHi-C (right) data. Solid lines indicate expected counts distributions, according to the CHiCAGO background model. The expected distribution was derived by taking samples from the null model, then kernel smoothing the resultant histogram to form a curve for visualisation purposes.

In addition to Brownian collisions, background in CHi-C is generated by assay artefacts, such as sequencing errors. We model this “technical noise” component as a Poisson random variable whose mean depends on the properties of interacting fragments, but is independent of genomic distance between them.

We further assume that these two sources of background counts are independent. Therefore, the combined background distribution can be obtained as a convolution of negative binomial (Brownian collisions) and Poisson (technical noise) distributions that is known as the Delaporte distribution.

We first construct this null distribution from the data in a robust way, based on all possible fragment pairs (including those that have zero observed read counts). We then find the pairs with counts that greatly exceed the expected background level (**Fig. 2**; as described in the next section). The full mathematical specification of the algorithm is given in **Additional file 1**.

#### Background estimation in asymmetrical interaction matrices

A practical advantage of the two-component background model is that the Brownian and technical normalisation factors can be estimated on separate subsets of data, each of which predominantly represents only one background component.

The dependence of background levels on the distance between fragments is particularly apparent at relatively short genomic distances (up to ~1-2Mb), where the observed read counts considerably exceed those observed at longer ranges and for trans-chromosomal interactions. Thus, within this range, counts arising from Brownian collisions largely dominate the technical noise, and hence the Brownian component can be estimated while ignoring the technical noise. By borrowing information across all interactions in this distance range, we can infer Brownian component parameters precisely (**Fig. 4**, **Suppl. Fig 1 in Additional file 2**). We follow Imakaev *et al*. [8] in assuming that fragment-level biases have a multiplicative effect on the expected read counts for each fragment pair. However we estimate “bait-specific” and “other end-specific” bias factors differently, accounting for the asymmetry of CHi-C interaction matrices.

**Figure 4:**
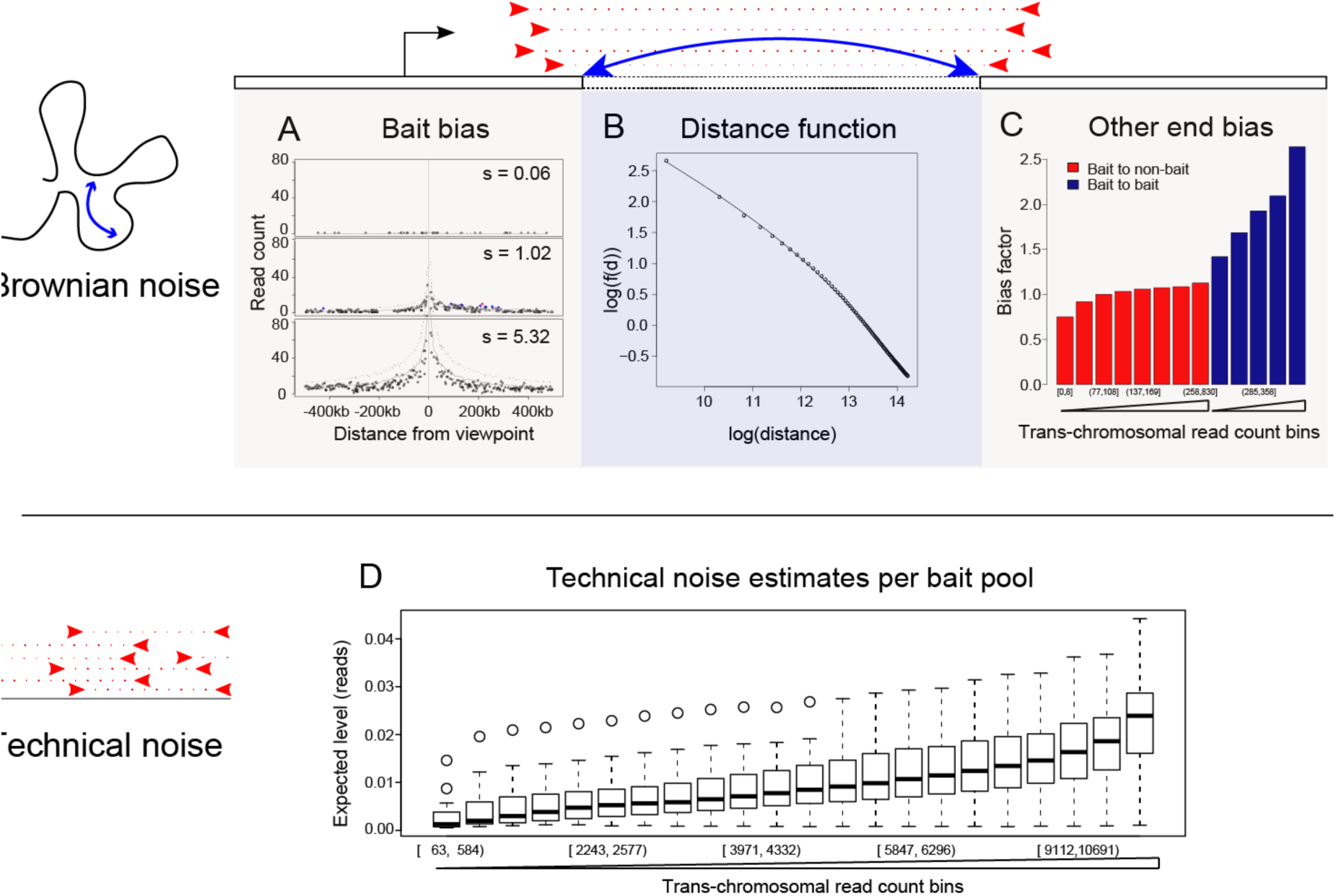
Sources of background counts and bias accounted for by the CHiCAGO model, illustrated with GM12878 data. (A-C) represent different factors that the Brownian background component models: (A) multiplicative bait-specific bias (shown are three representative distance profiles for three different values of the bait-specific bias factor); (B) distance dependency, plotted on a log-log scale; (C) multiplicative other-end bias (each bar represents a pool of other ends defined by a range of trans-chromosomal read pairs accumulated by each other end; bait-to-bait interactions are pooled separately). (D) Technical noise is estimated separately for each combination of bait and other-end pools, with pool membership defined by the number of accumulated trans-chromosomal read pairs. Here, we show the technical noise level estimates for each bait-other end pool combination, grouped together by bait pool.

The bait-specific factors reflect the technical biases of both Hi-C and sequence capture, as well as local effects such as chromatin accessibility. We estimate these factors in a way that is robust to the presence of a small fraction of interactions in the data. **Fig. 4A** provides examples of three baits with very diverse bias factors, illustrating that local read enrichment correlates with the bias factor.

Estimating other end-specific bias factors poses a challenge, as the majority of interactions are removed at the capture stage that enriches for only a small subset of interactions with baits. We assume that the overall fragment-level read count corresponding to *trans*-chromosomal pairs primarily reflects the general “noisiness” of a fragment (a similar approach has been taken independently in Dryden *et al*. [6]). While we do not preclude the presence of individual trans-chromosomal interaction signals, our reasoning that the overall per-fragment levels of trans-chromosomal pairs are dominated by noise is supported by evidence from Hi-C and random ligation control data (**Suppl. Fig 3 In Additional file 2**). We therefore pool fragments according to this property and estimate bias factors for each pool. As expected, bias factors are higher for fragments associated with higher numbers of trans-chromosomal read pairs (**Fig. 4C**). Similarly, baits detected at the “other ends” of bait-to-bait pairs had higher background levels than non-baits, as expected given the preferential recovery of “double-baited” ligation products at the capture stage.

In parallel, we compute the dependence between the Brownian background component and linear chromosomal distance (plotted in **Fig. 4B** for GM12878 CHi-C data). It can be seen that this dependence approximately follows a piecewise power law, consistent with previous studies on the subject, both theoretical and experimental [21, 22]. We further show by cross-validation that the estimate of this dependence is stable (**Suppl. Fig 2 In Additional file 2**), and therefore unlikely to be influenced by bait-specific or interaction-specific signals.

To estimate the magnitude of technical noise, we again use the per-fragment total trans-chromosomal read pairs (see Methods). In doing so, we assume that the contribution of true signals from specific trans-chromosomal looping interactions, as well as from Brownian collisions between chromosomes to the total trans-chromosomal counts, is negligible for reasons outlined above (**Suppl. Fig. 3 in Additional file 2**). Indeed, as we see in **Fig. 4D**, the expected level of technical noise is typically a small fraction of a count.

The estimated parameters of both background components are then combined into the Delaporte distribution. In **Suppl. Fig 4 (Additional file 2)**, we show evidence that CHiCAGO’s parameter estimation procedures are robust in the presence of undersampling; the implications of undersampling in CHi-C data are further examined in the Discussion.

After appropriate normalisation and bias correction, we detect fragment pairs showing read coverage higher than expected under the Delaporte assumptions with a one-tailed hypothesis test.

#### Weighted multiple testing correction for Capture HiC

For a typical mammalian genome, we test billions of hypotheses – one for each possible bait-other end pair. As a result, the p-values must be corrected to account for multiple testing. Standard multiple testing procedures assume that interactions are equally likely at all distances. However, in CHi-C data, we perform far more tests to verify the significance of interactions at large distances, where we would expect considerably fewer true interaction events. Consistent with this, the use of a single p-value threshold leads to results that consist mostly of erroneous distal and trans-chromosomal counts (**Fig. 5B**–**C**).

**Figure 5:**
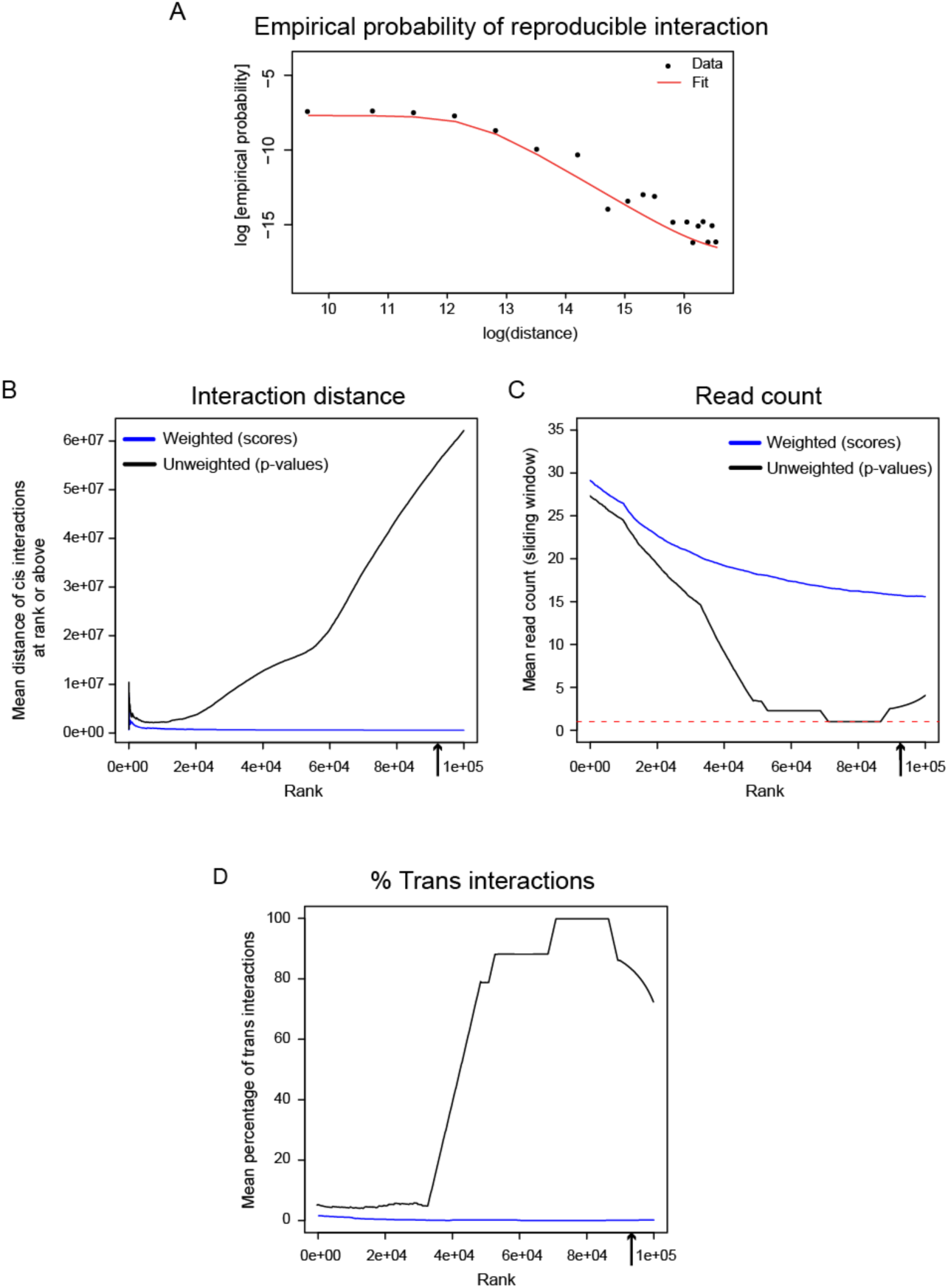
CHiCAGO multiple testing approach schematic. (A) Empirical probability of reproducible interaction (used to generate weight profiles) as a function of interaction distance, generated on two replicates of GM12878 cells, assessed for the 100,000 top-scoring interactions. (B-D) The effects of applying p-value weighting to the GM12878 data. The arrow on the x-axis indicates the number of significant interactions called in the weighted data. Upon applying weighting we see a decrease in the interaction distance amongst cis-interactions (B). Pvalue weighting increases the mean read count of called interactions (C) and decreases the prevalence of transchromosomal interactions (D).

To address this issue, the long-range and trans-chromosomal interaction tests need to be more stringent than the short-range ones. We achieve this with an approach based on p-value weighting [15, 23]. This procedure permits a smooth change of behaviour with distance, thereby bypassing the need to choose a hard distance threshold. Briefly, we assign each fragment pair a weight, estimating how probable it is that the fragments interact. The weights are then used to adjust the p-values (see **Additional file 1** for full specification). P-value weighting can be seen as a simplified version of the empirical Bayesian treatment, with weights related to prior probabilities. One practical advantage of this method for our framework is that it avoids the need to make specific assumptions about the read count distribution of true interactions, which would be required for computing Bayes factors.

The optimal choice of weights depends on the relative abundance of true positives at each bait–other end distance. We estimate this abundance by assessing reproducibility across samples and fitting a bounded logistic curve to the observed reproducibility levels at different distances. Generally similar weight profiles were obtained in GM12878 and mESC cells, and swapping them between these two datasets yielded highly correlated score profiles (**Fig. 5A** and **Suppl. Fig. 5 in Additional file 2**). This is consistent with our expectation that weights are largely independent of specific cell type and organism given comparable genome sizes, as they predominantly reflect the overall distance distribution of true interactions. The emerging multi-replicate CHi-C datasets will further refine our weight estimates and assess their dependence on the particulars of the model system.

We illustrate the impact of the weighting procedure on GM12878 and mESC CHi-C data by comparing the properties of the 100,000 top-scoring interactions, called either with or without weighting. The reproducibility of interaction calls decreases with bait–other end distance (**Fig. 5A** and **Suppl. Fig. 5A in Additional file 2**). As a result, the “weighted” significant interactions generally span a much shorter range than the unweighted ones (**Fig. 5B** and **Suppl. Fig. 5B in Additional file 2**). This is consistent with the biological expectation that promoter-interacting regions, such as enhancers, are enriched in the relative vicinity of their targets. Another consequence of the weighting procedure is that the average read count is much higher in the weighted calls (**Fig. 5C** and **Suppl. Fig. 5C in Additional file 2**). Strikingly, many of the unweighted calls are based on only one read pair per interaction. As the vast majority of fragment pairs attract no reads at all, low p-values for single-read-pair interactions are expected. However, due to the very large number of possible fragment pairs (approximately 18.5 billion in both the GM12878 and the mESC data), we still expect thousands of single-read-count calls to be generated by technical noise. These spurious calls, the majority of which correspond to trans-chromosomal pairs (**Fig. 5D** and **Suppl. Fig. 5D in Additional file 2**), are generally non-reproducible and are therefore excluded by the weighting procedure.

In conclusion, the p-value weighting procedure implemented in CHiCAGO provides a multiple testing treatment that accounts for the differences in true positive rates at different bait–other end distances, thus improving the reproducibility of interaction calls.

### Promoter interactions detected by CHiCAGO: validation and key properties

We validated CHiCAGO by assessing the functional properties of significant interactions detected with it in human GM12878 [3] and mouse ES cells [4]. **Table 1** displays summary statistics for each sample, showing the generally similar numbers of detected significant interactions, both overall and per bait, despite the differences in the organism and cell type between them.

**Table 1:** The properties of CHiCAGO-detected interactions in GM12878 human LCLs and mouse ES cells (mESC)

|  | GM12878 | mESC |
| --- | --- | --- |
| Number of captured baits | 22076 | 22459 |
| Total number of unique captured read pairs | Rep 1: 46542745<br>Rep 2: 118813226<br>Rep 3: 73881698 | Rep 1: 59963697<br>Rep 2: 82026534 |
| Number of significant interactions | 92457 | 81459 |
| Mean number of significant interactions per bait | 4.19 | 3.63 |
| Median distance of cis-chromosomal interactions | 173926 bp | 255330 bp |

#### Enrichment for regulatory features

We first assessed the enrichment of promoter-interacting fragments for histone marks associated with active (H3K4me1, H3K4me3, H3K27ac) and repressed (H3K27me3, H3K9me3) chromatin, as well as for the binding sites of CTCF, a protein with a well-established role in shaping nuclear architecture [24]. To this end, we compared the observed and expected numbers of significant other ends overlapping with these features. To estimate the expected degree of overlap, we drew multiple permutations of the promoter-other end pairs not detected as interacting, such that the overall distribution of their spanned distances matched the distribution for the true interactions.

**Fig. 6** shows the observed and expected numbers of CHiCAGO other ends (yellow and blue bars, respectively) that overlap with the regulatory features in GM12878 and mESCs (panels A and B, respectively; 95% confidence intervals are shown as error bars). Consistent enrichments over expected values were found for active histone marks (H3K4me1, H3K4me3, H3K27ac) in both cell types, in line with the expectation that looping interactions preferentially link promoters and remote regulatory regions such as enhancers. We also found that promoter-interacting fragments were strongly enriched for CTCF binding sites, as previously reported [9, 24]. Interestingly, promoter-interacting fragments were also enriched for repressed chromatin marks, in particular for H3K27me3 in mESCs, supporting the role of Polycomb in shaping nuclear architecture in this cell type [5].

**Figure 6:**
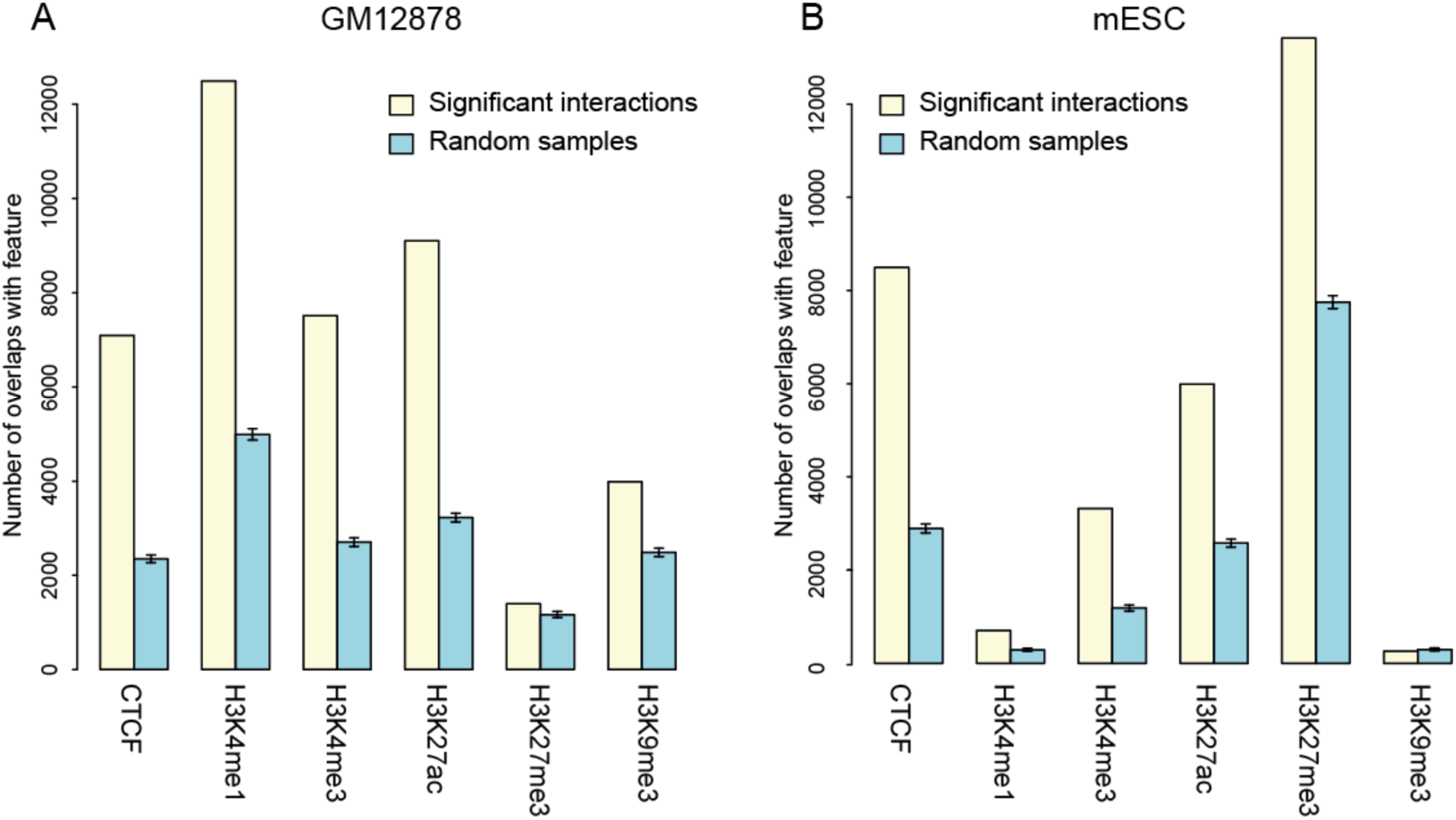
Chromatin features of promoter-interacting fragments detected using CHiCAGO. Yellow bars indicate overlaps with cis-interacting fragments within 1Mb from promoter baits; blue bars indicate expected overlap values based on 100 random subsets of *Hindlll* fragments. These subsets were selected to have a similar distribution of distances from gene promoters as the interacting fragments. (A) GM12878 CHi-C data. Chromatin features are obtained from the ENCODE project [25]; (B) mESC CHi-C data. Chromatin features are obtained from the mouse ENCODE project [26]. These plots are generated automatically by the CHiCAGO pipeline.

Assessing the enrichment of promoter-interacting fragments for known regulatory features can serve as a useful quality control for CHi-C samples. To this end, CHiCAGO automatically generates enrichment barplots similar to **Fig. 6** for each sample, integrating interaction calls with user-specified genomic annotations, such as ChIP-seq peaks.

#### Enrichment for GWAS SNPs

The majority of disease-associated SNPs identified in genome-wide association studies (GWAS) localise to non-coding regulatory regions, away from annotated promoters, posing a significant challenge in identifying their putative target genes [27]. We asked whether promoter-interacting regions detected by CHiCAGO in human cells are enriched for GWAS SNPs, which would potentially reflect their presence in long-range regulatory sequences and thus suggest a putative functional role in disease.

We assessed the enrichment of promoter-interacting regions in GM12878 cells for sets of GWAS catalogue SNPs from Maurano *et al*. [27]. These sets reflect the grouping of GWAS traits into broader categories, such as autoimmune disease (AI), neurological/behavioural traits (NB) and kidney/liver/lung disorders (KLL). We used the software package GOShifter (Genomic annOtation Shifter, [28]) that infers the significance of overlap by locally shifting genomic annotations (in our case, the “other ends” of CHiCAGO-detected promoter interactions), thus reducing the effect of genomic biases and LD structure. We observed a significant enrichment of CHiCAGO “other ends” for SNPs associated with autoimmune diseases (GOShifter p=0.001), but not with neurological/behavioural traits (p=0.801) or kidney/liver/lung disorders (p=0.876). This selective enrichment for autoimmune SNPs is consistent with GM12878 being a lymphocyte-derived cell line and replicates the original findings of Mifsud *et al*. [3].

We further confirmed that the enrichment for AI disease-associated SNPs was specific to promoter-interacting fragments. We used the same approach as in the previous section to generate 100 random samples of distance-matched “negative” (non-significant) interactions and tested the other ends of these interactions forSNP enrichment. The enrichment for Al-associated SNPs was selectively observed in the “true”, but not in the “negative” set, and neither set was enriched for the NB- and KLL-associated SNPs (**Fig. 7**).

**Figure 7:**
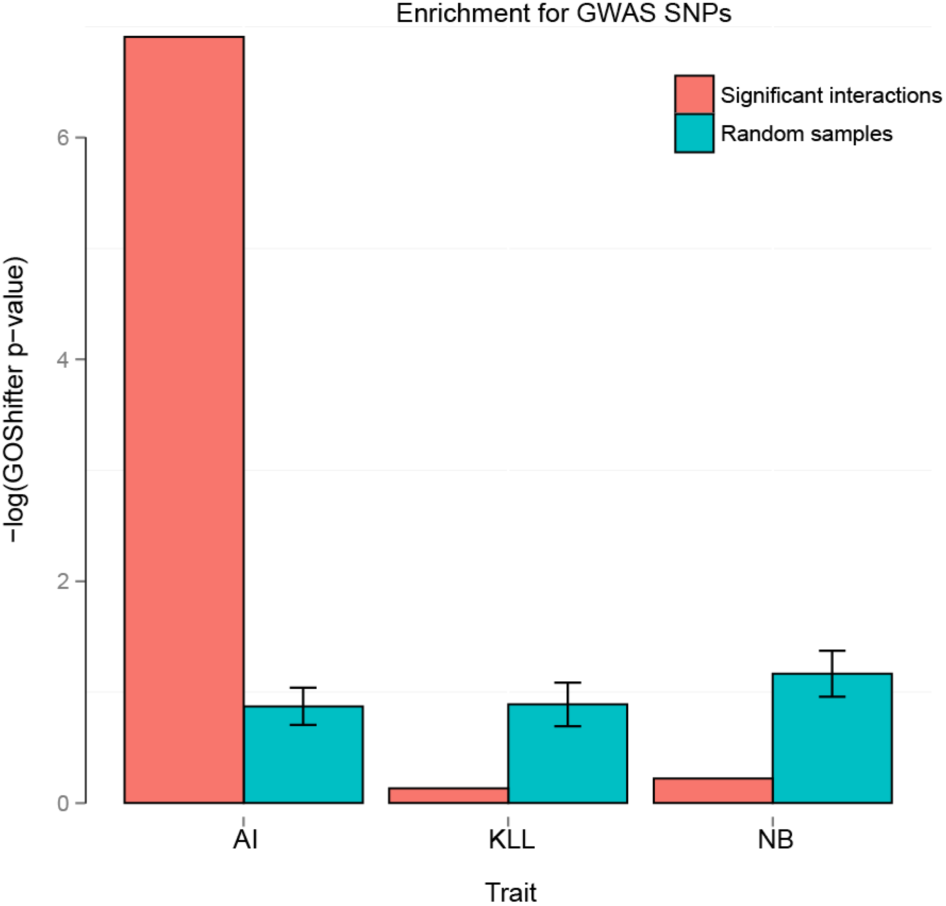
Significant enricliment for GWAS SNPs at CHiCAGO-detected interactions in liuman iymplioblastoid cells. Enrichment for SNPs associated with autoimmune immune diseases (AI), kidney/liver/lung (KLL) and neurological behaviour (NB) disorders [27] in the CHiCAGO-detected interactions in the GM12878 cell line. The barplot shows p-values for the enrichment of each disorder; red bars indicate p-values computed in interacting-fragments; blue bars indicate p-values computed in 100 random subsets of *Hindlll* fragments selected to have a similar distribution of distances from gene promoters as the interacting fragments. This analysis was performed using the software package GoShifter (Genomic Annotation Shifter, [28]).

Taken together, these results demonstrate the power of using CHi-C data to link GWAS SNPs with their putative target genes in a cell-type-specific and high-throughput manner. We expect this to be one of the key applications of CHi-C in future clinical studies.

#### Capability to drive transgene expression in vivo

TRIP (Thousands of Reporters Integrated in Parallel) is a novel experimental technique to assess the influence of local chromatin context on gene expression. In TRIP analysis, a barcoded transgene reporter is randomly integrated into thousands of genomic locations in parallel, and the transcriptional activity at each location is then monitored. Here we integrated the published TRIP analysis dataset in mESCs [29] with the CHiCAGO mESC calls [4], comparing the transcriptional activity at promoter-interacting regions with the activity elsewhere, over a range of genomic distances.

Consistent with the observation from the original TRIP study, we found that the distance from the nearest promoter was a strong determinant of transgene expression levels (**Fig. 8**). However, transgenes mapping to promoter-interacting fragments consistently showed higher expression levels across the whole range of genomic distances, as confirmed by linear regression (odds ratio=2.27; Wald test p<0.001). This result provides functional evidence that CHiCAGO-detected promoter-interacting fragments preferentially possess transcriptional regulatory activity.

**Figure 8:**
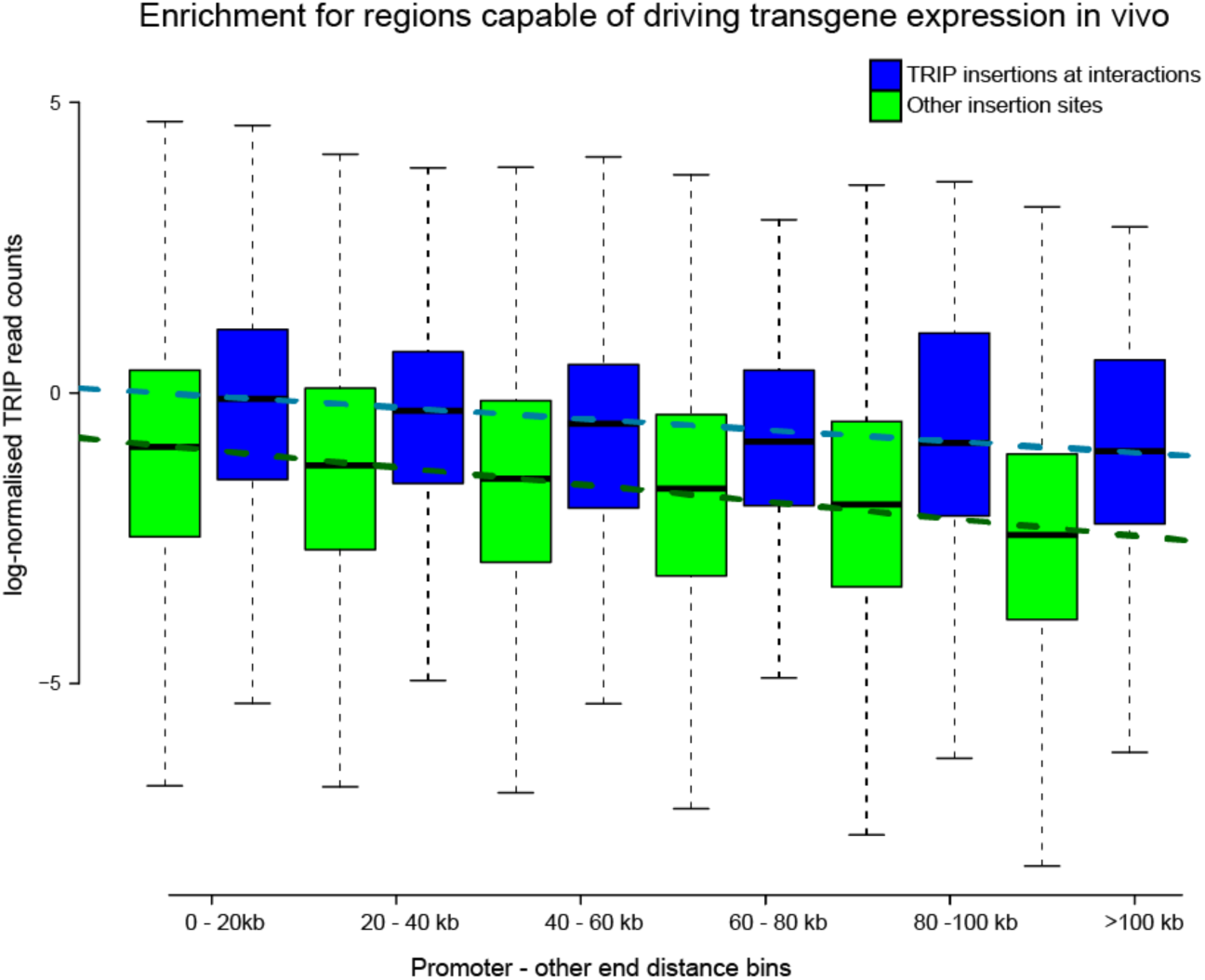
Enrichment of promoter-interacting fragments for regions capable of driving transgene expression in mESCs. TRIP (Thousands of Reporters Integrated in Parallel) assesses the influence of local chromatin context on gene expression. This is achieved by integrating a barcoded transgene reporter into thousands of genomic locations in parallel and monitoring the transcriptional activity at each location [29]. Normalised RNA read counts from reporter insertions are separated according to (i) their overlap with Hindlll fragments engaging or not in interactions; (ii) their promoter-other end distance. For non-interacting Hindlll fragments, distance is measured from the nearest promoter in the linear sequence. Blue and green boxplots indicate read count summary statistics for promoter-interacting and non-interacting *Hindlll* fragments, respectively. Each dashed line shows the regression of median log-normalised read counts against promoter-other end distance bin, considering promoter-interacting (blue) and non-interacting (green) *Hindlll* fragments separately.

#### Promoter-promoter networks

Interactions where both fragment ends are baited (referred to as bait-to-bait interactions) represent contacts between gene promoters. These interactions are of special interest because they may help to identify sets of co-regulated genes recruited to either shared transcription factories [30] or repression networks such as those mediated by Polycomb proteins [5].

As an illustration of CHiCAGO’S potential in identifying sets of co-regulated genes, we show CHiCAGO-detected bait-to-bait interactions involving histone promoters present on chromosome 6 in GM12878 cells (**Fig. 9**). We see that histone promoters frequently interact with other histone promoters, more so than with promoters of other genes in the same genomic region, consistent with previous observations [4, 31, 32].

**Figure 9:**
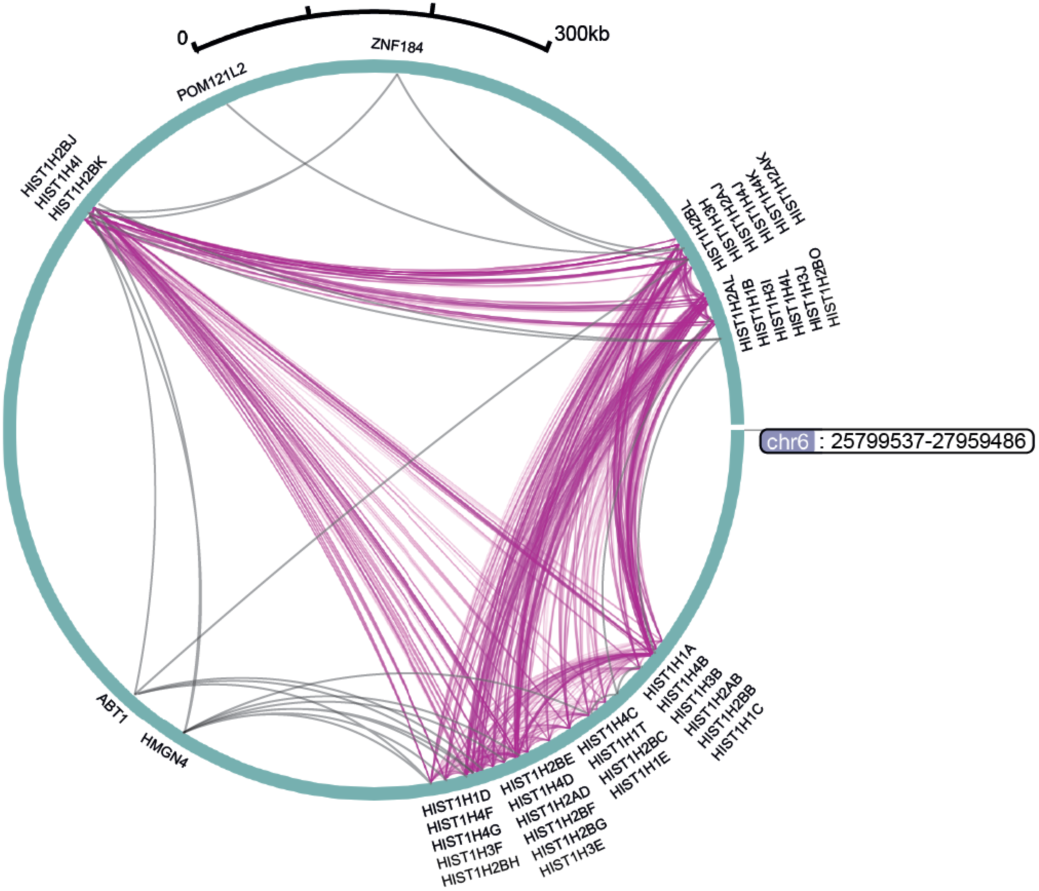
Circlet view of promoter-promoter interactions for histone genes in GM12878. Interactions where histone gene promoters are engaged at both fragment ends are shown in dark magenta. Interactions where histone gene promoters are interacting with non-histone gene promoters are shown in grey. The WashU EpiGenome Browser [16, 17] was used to create this figure.

#### Extremely long-range promoter interactions map within broader Hi-C contact regions

We took advantage of the pre-capture Hi-C dataset in mESCs [4] from to compare CHiCAGO-detected interactions in Promoter CHi-C with the broader-scale interaction signals detectable in Hi-C. The Promoter CHi-C dataset has over 10-fold higher coverage at promoters compared to the respective Hi-C sample [4], and thus we would expect a corresponding increase in the sensitivity of detecting promoter-containing interactions. Consistent with this, while some stronger interactions in the short range (<1Mb) could be visually distinguished on Hi-C interaction matrices (**Fig. 10A**), more than 80% of CHi-C interactions in this range localised away from Hi-C interacting regions detected with HOMER [33] at a 25kb resolution (**Fig. 10B**). In contrast, we found that more than 80% extremely long-range (>10Mb) cis-chromosomal interactions and 45% trans-chromosomal interactions mapped within the broader (1Mb-wide) Hi-C contact areas (**Fig. 10C**). However, only a small minority of these megabase-scale contact areas contained CHi-C interactions (~3% of >10Mb cis-chromosomal and ~0.5% trans-chromosomal, as illustrated in **Fig. 10D** and **Suppl. Fig. 6 in Additional file 2**). Taken together, these results are consistent with a high specificity and resolution of CHiCAGO long-range interaction calling. At the same time, they warrant a further examination of the relationship between specific looping interactions and higher-order chromosomal contacts.

**Figure 10:**
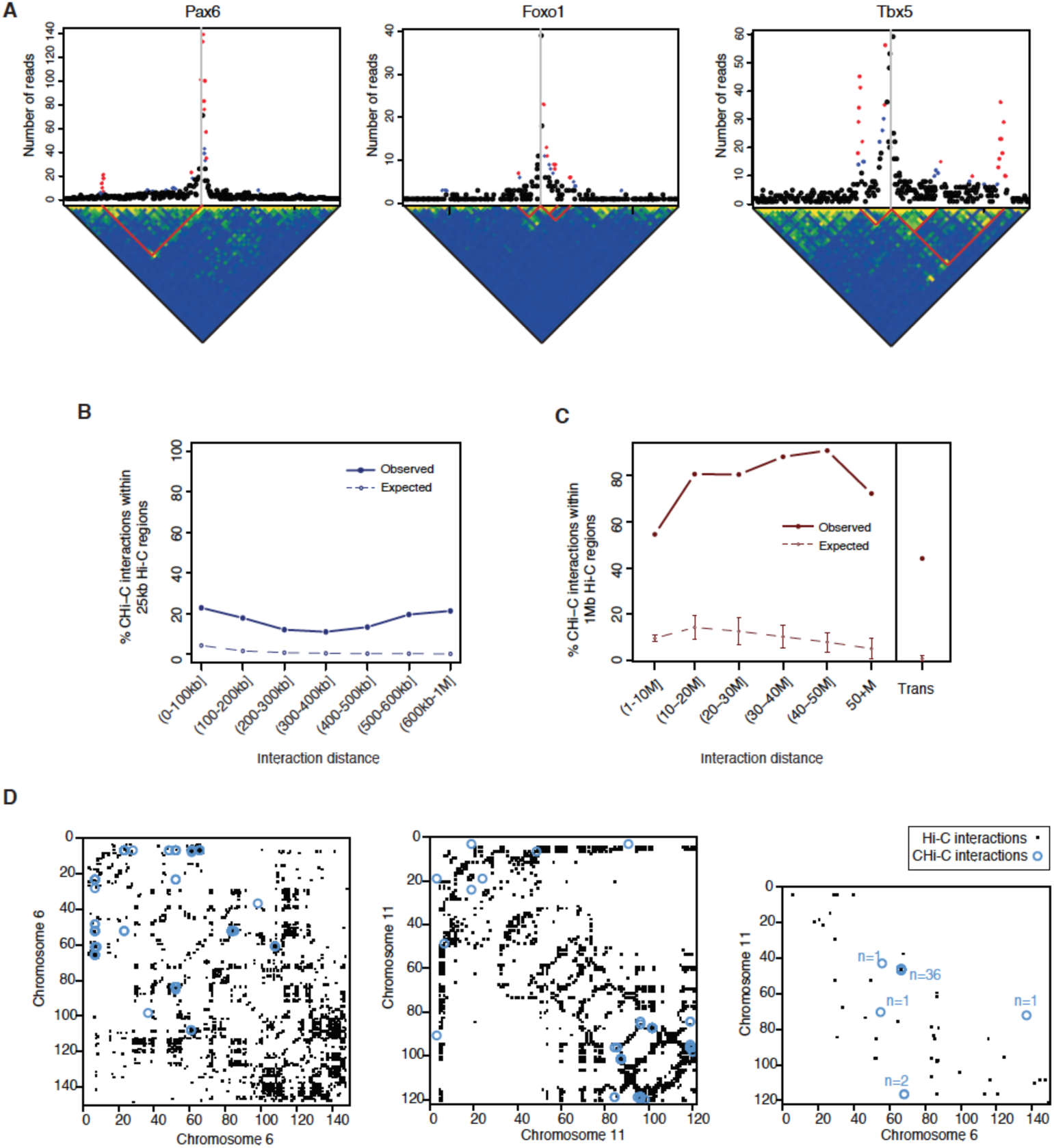
Comparison of interactions detected in CHi-C and Hi-C data. **(A)** Top panels: Plots showing the read counts from bait-other end pairs within 750kb (upstream and downstream) of 3 baits, containing the *Pax6, Foxo1* and *Tbx5* promoters (from left to right). Significant interactions detected by CHiCAGO (score >= 5) are shown in red, and sub-threshold interactions (3 <= score < 5) are shown in blue. Bottom panels: raw Hi-C matrices at 25-kb bin resolution within the corresponding 1.5Mb regions. The bottom corners of the red lines indicate example bin pairs, within which significant interactions were detected in the CHi-C data. **(B)** Mapping of short-range (<1Mb) CHi-C interactions within 25-kb interacting bins detected in the Hi-C data. Filled circles show the observed fraction of CHi-C interactions mapping within the Hi-C interacting bins; open circles show the expected fraction estimated by a permutation strategy accounting for genomic structure (see Methods for details). The standard deviations across 100 permutations are not shown as they are smaller than point size. **(C)** Mapping of long-range (>1Mb) CHi-C interactions within 1-Mb interacting bins detected in the Hi-C data. Filled circles show the observed fraction of long-range *cis*- and *trans*-chromosomal interactions detected in the CHi-C data that map within the Hi-C interacting bins. Open circles show the expected fraction estimated by a permutation strategy accounting for genomic structure (see Methods for details). Error bars show standard deviation across 100 permutations. **(D)** The overlap of long-range (>5Mb) interacting fragment pairs detected in CHi-C data (blue circles) and interacting 1Mb bin pairs detected in the Hi-C data (black squares) on chromosomes 6 (left) and 11 (centre), and for *trans*-interactions between these chromosomes (right). All panels (A)-(D) present mouse precapture ES-cell Hi-C data from [4].

## DISCUSSION

In this paper, we presented the CHiCAGO algorithm for Capture Hi-C analysis and demonstrated its efficacy in detecting interactions enriched for regulatory chromatin features and relevant GWAS SNPs.

Our approach is based on the assumption that “significant” interactions emerge as outliers on a distance-dependent local background profile. This assumption is shared by most other tools for interaction detection in 3C-like data and seems reasonable for the purposes of identifying regulatory interactions. Indeed, it can be expected that regulatory events such as transcription factor binding will stabilise the chromatin loop, leading to interaction frequencies or retention times beyond those generated by random collisions due to Brownian motion.

This expectation is supported by the observation that CHiCAGO-detected interactions are selectively enriched for regulatory chromatin features, even when located in regions with high background interaction levels.

While the conceptual interpretation of “significant” interactions is shared between CHiCAGO and algorithms developed for other types of 4C and Hi-C data, there are key differences in terms of the underlying background model, the normalisation strategy and the multiple testing procedure.

Existing tools model Hi-C background with a broad range of distributions, both discrete (binomial [19, 34], negative binomial [6]) and continuous (Weibull [7, 9], normal [13]). In CHiCAGO, we instead opted for a two-component convolution model that incorporates two count distributions: a negative binomial and a Poisson. In doing so, we were motivated by the fact that distance-dependent Brownian collisions and technical variability are two distinct background count-generating processes, whose properties are best learned separately on different subsets of data. Indeed, signals from Brownian collisions ostensibly dominate the background at short distances, to the extent that technical variability is barely detectable. In contrast, at large linear distances between fragments, Brownian collisions are too weak for their count distribution to be estimated directly. Thus, we infer this distribution by extrapolation.

Borrowing information across baits to learn the background model, as CHiCAGO does, requires careful normalisation across interactions. While Hi-C background depends on a number of known parameters, such as fragment length and GC content [10], we, along with others [7, 8, 35], have opted to avoid any specific assumptions about noise structure, particularly given the increased complexity and asymmetric nature of capture Hi-C noise compared with conventional Hi-C. Assuming that interactions are subject to multiplicative bait- and other-end-specific bias, as we did in learning the Brownian background component, parallels the assumptions of the Hi-C iterative correction approach by Imakaev *et al*. [8] and is generally consistent with data from molecular dynamics simulations of chromatin fibres [21]. In modelling technical noise, we assumed it to be reflected in the numbers of *trans*-chromosomal interactions involving the same fragment. A similar strategy has been applied independently in a recently published Capture Hi-C study [6]; the same authors also proposed an iterative correction algorithm for Capture Hi-C data [7] (software not publicly released) that may complement the approaches taken here.

Multiple testing issues are important in genomic analyses and, in attempting to address these issues, a number of bespoke approaches have been developed [23, 36]. The specific challenge of multiple testing in Hi-C data is that we expect the fractions of true positives to vary depending on the genomic distance between the fragments; in fact, the majority of tests are performed with interactions spanning large distances or spanning different chromosomes, where true positive signals are least expected. CHiCAGO’s multiple testing procedure is based on the p-value weighting approach by Genovese *et al*. [15], which is a generalisation of a segment-wise weighting procedure by Sun *et al*. [37]. These approaches have been used successfully to incorporate prior knowledge in genome-wide association studies [38–40] and are emerging in functional genomics analyses [41, 42]. In using the reproducibility of significant calls across replicates as an estimate of the relative true positive rate, we have taken inspiration from the irreproducible discovery rate (IDR) approach [43] used to determine peak signal thresholds in other types of genomics data, such as ChIP-seq.

Note that, in this setting, IDetermining Transcription FactorsDR cannot be used verbatim for choosing signal thresholds, as the relationship between Capture Hi-C signal and reproducibility does not satisfy IDR assumptions, likely because of undersampling issues (not shown). Importantly, conventional false discovery rate (FDR-) based approaches for multiple testing correction [44] are unsuitable for these data. Indeed, CHi-C observations (read-pair counts) are discrete and many of them are equal to either zero or one. This leads to a highly non-uniform distribution of p-values under the null, violating the basic assumption of conventional FDR approaches. The “soft-thresholding” approach used in CHiCAGO shifts the -log-weighted p-values such that non-zero scores correspond to observations, where the evidence for an interaction exceeds that for a pair of near-adjacent fragments with no reads. More robust thresholds can then be chosen based on custom criteria, such as maximising enrichment of promoter-interacting fragments for chromatin features (**Fig. 6**; a user-friendly function for this analysis is provided as part of the Chicago R package - see the package vignette provided as **Additional file 3**). Based on this approach, we chose a signal threshold of 5 for our own analyses.

The undersampled nature of CHi-C data (particularly at longer distance ranges), although robustly handled by CHiCAGO, may lead to significant sensitivity issues when using thresholded interaction calls in comparative analyses. We therefore suggest performing comparisons based on the continuous score range. Potentially, differential analysis algorithms for sequencing data (such as DESeq2 [45]) may also be used to formally compare the enrichment at CHiCAGO-detected interactions between conditions at the count level, although power will generally be a limiting factor. As undersampling drives down the observed overlap of interactions called on different samples (**Suppl. Fig 4C** in **Additional file 2**), methods such as [46, 47] may be considered for formally ascertaining the consistency between datasets. Additional filtering based on the mean number of reads per detected interaction (e.g., removing calls with N<10 reads) will also reduce the impact of undersampling on the observed overlap, but at the cost of decreasing the power to detect longer-range interactions.

The p-value weighting approach used here is similar in spirit to an empirical Bayesian treatment, with the p-value weights related, but not identical, to prior probabilities. Bayesian approaches are widely used (including, recently, for signal detection in conventional Hi-C [48]), and the Bayes factors and posterior probabilities they generate are potentially more intuitive than weighted p-values. However, the p-value weighting approach used here has the advantage of not making any specific assumptions of the read distribution of “true interactions”, beyond their having a larger mean. Both approaches open the opportunity of incorporating prior knowledge, beyond the dependence of reproducibility on distance - for example, taking into account the boundaries of topologically associated domains (TADs [49]), higher-order contact domains and chromosomal territories. We choose not to do this currently, because the exact relationship between these genomic properties and looping interactions still requires further investigation, and incorporating these relationships *a priori* prevents their investigation in post-hoc analyses. Active research in this area makes it likely that much more will be known about the determinants of loop formation in the near future, enabling a more extensive use of prior knowledge in interaction detection, potentially with a formal Bayesian treatment.

The downstream analyses of CHiCAGO results provided in this paper confirm the enrichment of promoter-interacting regions for regulatory features and disease-associated variants. These results demonstrate the enormous potential of Capture Hi-C for both functional genomics and population genetics, and this assay will likely be applied in multitudes of other cell types in the near future. Therefore, user-friendly, open-source software for robust signal detection in these challenging data will be a welcome addition to the toolkits of many bioinformaticians and experimentalists alike. We have developed CHiCAGO with the view of addressing this need. Furthermore, we expect the statistical foundations of CHiCAGO, particularly the convolution background model and the multiple testing procedure, to be potentially useful in a broader range of Hi-C-related assays.

## CONCLUSIONS

The publicly available, open-source CHiCAGO pipeline presented here [50] produces robust and interpretable interaction calls in Capture Hi-C data. Promoter-interacting fragments identified using this algorithm are enriched for active chromatin features, GWAS SNPs and regions capable of driving transgene expression, indicative of regulatory looping interactions. While developed specifically for Capture Hi-C, the statistical principles of CHiCAGO are potentially applicable to other Hi-C-based methods.

### MATERIALS AND METHODS

#### Sample pre-processing

The publicly available HiCUP pipeline [51, 52] was used to process the raw sequencing reads. This pipeline was used to map the read pairs against the mouse (mm9) and human (hg19) genomes, to filter experimental artefacts (such as circularized reads and re-ligations), and to remove duplicate reads. For the CHi-C data, the resulting BAM files were processed into CHiCAGO input files, retaining only those read pairs that mapped, at least on one end, to a captured bait. The script bam2chicago.sh, used for this purpose, is available as part of the chicagoTools suite [50].

#### The CHiCAGO algorithm

A full description of the algorithm is given in **Additional file 1**. A tutorial on using the CHiCAGO package (the “vignette”) is provided in **Additional file 3**.

Briefly, to combine replicates, a “reference” replicate is created by taking the geometric mean of each fragment pair’s count across samples. Sample size factors are calculated by taking the mean ratio to the “reference” replicate, in a manner similar to the sample normalisation strategy implemented in DESeq [53]. Final counts are derived as the weighted sum of counts across replicates, where the weights are the sample size factors.

Background from Brownian collisions is assumed to have Negative Binomial distribution, with mean s_i_s_j_f(d_ij_) and dispersion r, where i indexes over other ends and j indexes over baits.

Estimation of s_i_, s_j_, f(d) and r is performed in “proximal bins” - by default, 20kb bins that span the first 1.5Mb around each bait.

The distance function f(d) is estimated as follows:

- For each bait, take all of the other ends in a distance bin to get a mean count for that bin.
- f(d) is estimated in a distance bin by taking the geometric mean of the bin counts at that distance, across all baits.
- To interpolate f(d) from these point estimates, we use a cubic fit on a log-log scale.
- Outside of this distance range, we extrapolate linearly, assuming continuity of f and its first derivative.

The bait-specific scaling factors, s_j_, are estimated by considering each mean bin count divided by f(d), then taking the median of this ratio, across all bins associated with a bait.

The other end-specific scaling factors, s_i_, are estimated similarly, but with the other ends pooled together (the pools are chosen such that their content ends have similar numbers of trans counts) so that there is enough information for a precise estimate. The dispersion, r, is estimated using standard maximum likelihood methods.

The technical noise is assumed to have Poisson distribution, with mean λ_ij_. λ_ij_ is estimated from trans counts - again, first pooling fragments by the number of trans counts they exhibit. Specifically, to estimate the technical noise level for a putative interaction between a bait in pool A and an other end in pool B, we count the number of interactions that span between pools A and B, and divide this by |A | |B |, the total number of bait-other end fragment pairs from those pools.

P-values are called with a Delaporte model, representing the sum of two variables: a Negative Binomial variable with mean s_i_s_j_f(d_ij_) and dispersion r, and a Poisson variable with mean λ_ij_. A four-parameter bounded logistic regression model is assumed for p-value weighting (see next section and **Additional file 1** for more information).

The final CHiCAGO score is obtained from soft-thresholding the -log(weighted p-value). Specifically, the score is max(-log(p) * log(w) -log(w_max_), 0). where w_max_ is the maximum attainable weight, corresponding to zero distance. For the downstream analyses in this paper, interactions with CHiCAGO scores >= 5 were considered as “significant interactions”.

#### P-value weighting parameter estimation

The p-value weighting function has four parameters: α, β, γ, and δ (full details are given in **Additional file 1**). We can estimate these parameters from a candidate data set, provided that it has multiple biological replicates, as follows. We split the data into subsets that contain approximately equal numbers of baits. (By default, 5 subsets are used.) The reproducible interactions are defined as those where the stringent threshold of log(p) < -10 is passed in all biological replicates. Now, for each subset, we take a series of genomic distance bins (with the default breaks occurring at 0, 31.25k, 62.5k, 125k, 250k, 500k, 1M, 2M, 3M, 4M,…, 16M), and we calculate the proportion of reproducible interactions out of the total number of possible interactions. The maximum likelihood estimates are calculated for each model parameter, using standard optimization methods [54]. Final parameter estimates are obtained by taking the median across the estimates from each subset. The two replicates of mESCs data [4] were used for estimating weights. For GM12878 [3], the first replicate was not used for weight estimation as it led to unstable estimation, likely due to the poorer quality of this replicate compared with the other two, consistent with its higher cis/trans read-pair ratios (data not shown). Recommendations on diagnosing unstable estimates are provided in the R package vignette (**Additional file 3**).

#### The Chicago R package

CHiCAGO was implemented as a package for the statistical environment R [55] taking advantage of the data.table objects [56] to optimise for both speed and memory. The fully-documented R package “Chicago” and the tutorial data package “PCHiCdata” are publicly available [50] and are part of Bioconductor [57]. A documented set of supplementary scripts (chicagoTools) for data pre- and post-processing and running Chicago in batch mode is also publicly available [50]. A typical Chicago job for two biological replicates of CHi-C data takes 2-3 h wall-clock time (including sample pre-processing from bam files) and uses 50G RAM. An example workflow in the form of an R package vignette is provided as **Additional file 3**. The description of free parameters and rationale for their settings is given in **Suppl. Table 1 (Additional file 2)**.

#### Assessment of feature enrichment

Enrichment for chromatin features at CHi-C interacting regions was assessed with respect to random *HindIII* fragments drawn in such a way as to match the distribution of the observed interaction distances. A 95% confidence interval for the expected overlap was obtained from 100 random draws. SNP enrichment at promoter interacting fragments was assessed using GOshifter [28].

#### Hi-C analyses

HOMER [33] was used on Hi-C data to compute binned coverage- and distance-related background in the Hi-C data and call significantly interacting bin pairs. Short-range *cis*-chromosomal interactions (<1Mb) were detected in 25-kb bins; long-range cis-chromosomal (>1Mb) and *trans*-chromosomal interactions were detected in 1Mb bins. Bin pairs with FDR-adjusted p<0.05 were considered significant. The significance of overlap between CHi-C promoter-interacting regions identified by CHiCAGO and the HOMER-detected interacting bin pairs in the Hi-C data was ascertained by permutation, while preserving the structural features of the data, as follows. *Cis*-chromosomal interactions were permuted across the baits while preserving the interaction distances. *Trans*-chromosomal interactions were permuted across chromosomes while preserving the relative chromosomal position of the interacting fragments.

#### Data access

Raw CHi-C, Hi-C and random ligation control data used in this study is available in ArrayExpress [58, 59] under accession numbers E-MTAB-2323 (GM12878) and E-MTAB-2414 (mESC), respectively. Capture design files, *HindIII* digest maps and CHiCAGO-detected significant interactions for GM12878 and mESC will be made publicly available prior to paper release.

## AUTHORS’ CONTRIBUTIONS

JC, PFP and MS designed the CHiCAGO algorithm; VP and DZ contributed statistical advice; JC, PFP, SWW and MS implemented the algorithm. SS, CO, BMJ and PF generated Capture Hi-C data and advised on their biological properties. PFP, CV, AD, JC and MS performed downstream validation analyses. JC, PFP and MS wrote the paper with critical input from all authors. MS supervised the work.

## ACKNOWLEDGEMENTS

The authors would like to thank Simon Andrews, Chris Wallace, Oliver Burren, and all members of the Spivakov, Fraser and Babraham Bioinformatics groups for helpful discussions. We are grateful to all our “wet-lab” collaborators (in particular, Mayra Furlan-Magaril, Mattia Frontini, Peter Rugg-Gunn and Willem Ouwehand) for using and testing CHiCAGO. This work has been funded by the Biotechnology and Biological Sciences Research Council and the Medical Research Council of the UK; DZ is funded by the European Molecular Biology Laboratory. Finally, we thank Laura Biggins for disambiguating the last two letters of CHiCAGO.

## ADDITIONAL FILES

Additional file 1: The mathematical specification of the CHiCAGO algorithm.

Additional file 2: Supplementary figures 1 through 6, and Supplementary Table 1.

Additional file 3: The CHiCAGO R package tutorial.

